# Evolution of influenza A virus hemagglutinin H1 and H3 across host species

**DOI:** 10.1101/2022.04.20.488870

**Authors:** Nídia S. Trovão, Sairah M. Khan, Philippe Lemey, Martha I. Nelson, Joshua L. Cherry

## Abstract

The Influenza A virus (IAV) hemagglutinin protein (HA) has been studied extensively, but its evolution has not been thoroughly compared among major host species. We compared H3 and H1 evolutionary rates among 49 lineages differentiated by host (avian, canine, equine, human, swine), phylogeny, and geography. Rates of nonsynonymous evolution, relative to synonymous rates, were higher in mammalian than avian hosts. Human seasonal HA and classical swine H1 accumulated 11-13 glycosylation sites, primarily in the antigenically important head domain, whereas lower numbers were maintained in other hosts. Canines had the highest ratio of nonsynonymous to synonymous changes in the more conserved stalk domain. Amino acid changes in canine viruses occurred disproportionately at residues located at protein interfaces. This suggests that they were adaptations affecting the major structural rearrangement of HA, which is critical for cell entry. These findings invite further study of how host ecology and physiology affect natural selection.

**AUTHOR SUMMARY:** Influenza virus evolution is of practical importance to health in addition to being an excellent system for the study of parasite/host evolution. Our work explores a largely untapped aspect of influenza evolution: sequence evolution in non-human hosts. This is important in its own right, in terms of both science and domestic animal health. It also puts the evolution of human influenza in a larger, comparative context. Our results also provide evidence concerning the evolution of the hemagglutinin stem domain, which has not been a focus of study but has new importance due to the development of stem-based universal influenza vaccines.

## INTRODUCTION

Influenza A and B viruses are the etiological agents of seasonal influenza in humans. Unlike influenza B viruses, which are only known to infect humans and seals, influenza A viruses of H3 and H1 subtypes infect humans and a multitude of non-human animals [1]. Influenza A viruses (IAVs) periodically jump between host species, presenting an ongoing pandemic threat [2, 3]. Sixteen different haemagglutinin (HA) subtypes in influenza A have been identified (H1 to H16), of which H1 and H3 are currently circulating in human populations [4]. Influenza A viruses of the H3 and H1 subtypes currently circulate in birds, humans, swine, dogs, and horses [5–11]. Human seasonal IAVs infect between 10% and 20% of the human population every year, causing an estimated 290 000 to 650 000 deaths annually [12]. IAVs also cause significant morbidity in animal populations, affecting swine and poultry production [13, 14] and the horse-racing industry [15–17].

Influenza viruses of the H1 and H3 subtypes are maintained in wild birds. During the twentieth century, spillover of avian viruses to humans caused the 1918 H1N1 [18] and 1968 H3N2 pandemics [19]. Avian viruses (H3N8) also became established in equines between the 1950s and 1960s [20]. Human viruses have transmitted dozens of times to pigs, establishing several independently evolving H1N1, H1N2, and H3N2 swine lineages on multiple continents [21]. In the early 2000s, two canine lineages emerged independently in the United States and in Asia through introductions from horses (H3N8) and birds (H3N2), respectively [8, 22]. More recently, a H1N1 virus jumped from swine to humans and caused the 2009 H1N1 pandemic [23]. The pandemic lineage quickly disseminated globally and transmitted back to swine repeatedly on multiple continents.

Mutations occur frequently during IAV replication by an error-prone RNA polymerase. The rate of synonymous substitution varies by host species, presumably due largely to differences in mutation rate [24]. This is affected by host-specific factors, including the number of viral replications per unit time and biochemical differences affecting replication fidelity. The rate of nonsynonymous evolution, relative to the mutation rate, is determined by the forces of natural selection. Nonsynonymous mutations, which change the amino acid encoded, are frequently deleterious, but may be adaptive. For example, they may diminish binding by existing antibodies [25] or improve binding to host cell receptors [26], particularly after a host switch.

Research on influenza evolution and adaptation has focused on the HA, the membrane glycoprotein present on the surface of the virus which is responsible for receptor binding and membrane fusion and which is under constant selection due to the host immune response.

Each subunit of the HA homotrimer consists of two domains. The immunodominant head domain is the principal target of antibody-mediated immunity and evolves more rapidly than the stalk domain, which is relatively conserved within and among subtypes [27–29]. Mutations in the HA head domain can help the virus evade antibodies, increasing its ability to infect individuals with immunity elicited by earlier strains. Some HA mutations create or eliminate glycosylation sites [30]. Although glycosylations can shield HA from antibodies, they can also interfere with receptor binding and are targets of innate immunity [31, 32]. Due to the different strengths of immune selection in different host species, the optimal number of glycosylations is expected to vary among them [33].

Influenza vaccine strains are updated twice a year to reflect antigenic changes in viruses circulating globally. However, due to limitations of the global surveillance and influenza vaccine manufacturing process, vaccine strains are periodically mismatched to viruses in circulation, leading to reduced vaccine effectiveness [34–36]. Vaccines are also available for swine [37, 38], canines [39], equines [40–42], and poultry [43], although in some cases updates of animal vaccine strains are not systematic and infrequent and their effectiveness and utilization is even lower than in humans [44, 45]. Studies of antigenic evolution have often relied on serological assays (hemagglutination inhibition (HI) or virus neutralization assays). However, they are not always well standardized or comparable across laboratories, and little antigenic data is available for non-human influenza. This impedes comparison of rates of protein and antigenic evolution of influenza in non-human hosts. In recent years, next-generation sequencing has vastly increased the volume of genetic data for studying evolution of HA in humans and other species.

Although vaccines have relied primarily on antibodies to the HA head for nearly a century, novel vaccine approaches may provide broader, longer lasting protection [46] by targeting, for instance, the HA stalk domain. The prospect of stalk-based vaccines gives a new importance to understanding evolution of the stalk.

Here, we compare the evolutionary dynamics of H1 and H3 head and stalk domains in humans, swine, birds, equines, and canines, including rates of synonymous and nonsynonymous change, the locations of changes in three-dimensional structures, and rates and patterns of gains and losses of glycosylation sites. Our results suggest that the head domain, which evolves rapidly in humans due to selection by the immune system, also does so in other mammals, though not to the same extent. We also find evidence of host-specific adaptations, especially in canines.

## RESULTS

### Sequence datasets

The final H3 dataset consisted of 873 sequences from viruses isolated between 1963 and 2019 from the following hosts: avian (n=178), human (n=101), swine (n=370), equine (n=87), and canine (n=137). For H1 there were 964 sequences, with collection dates between 1918 and 2009, from avian (n=136), human (n=186), and swine (n=642) hosts. The H3 sequences were divided into 16 lineages, and the H1 sequences into 33 (Figs. S1 and S2, and Files S1 and S2).

### Evolutionary Dynamics of Influenza A/H1 and A/H3 Across Host Species

Influenza A/H1 and A/H3 have frequently jumped from one host to another during the last century, establishing multiple geographic- and host-specific lineages (Fig. 1).

**Fig. 1:**
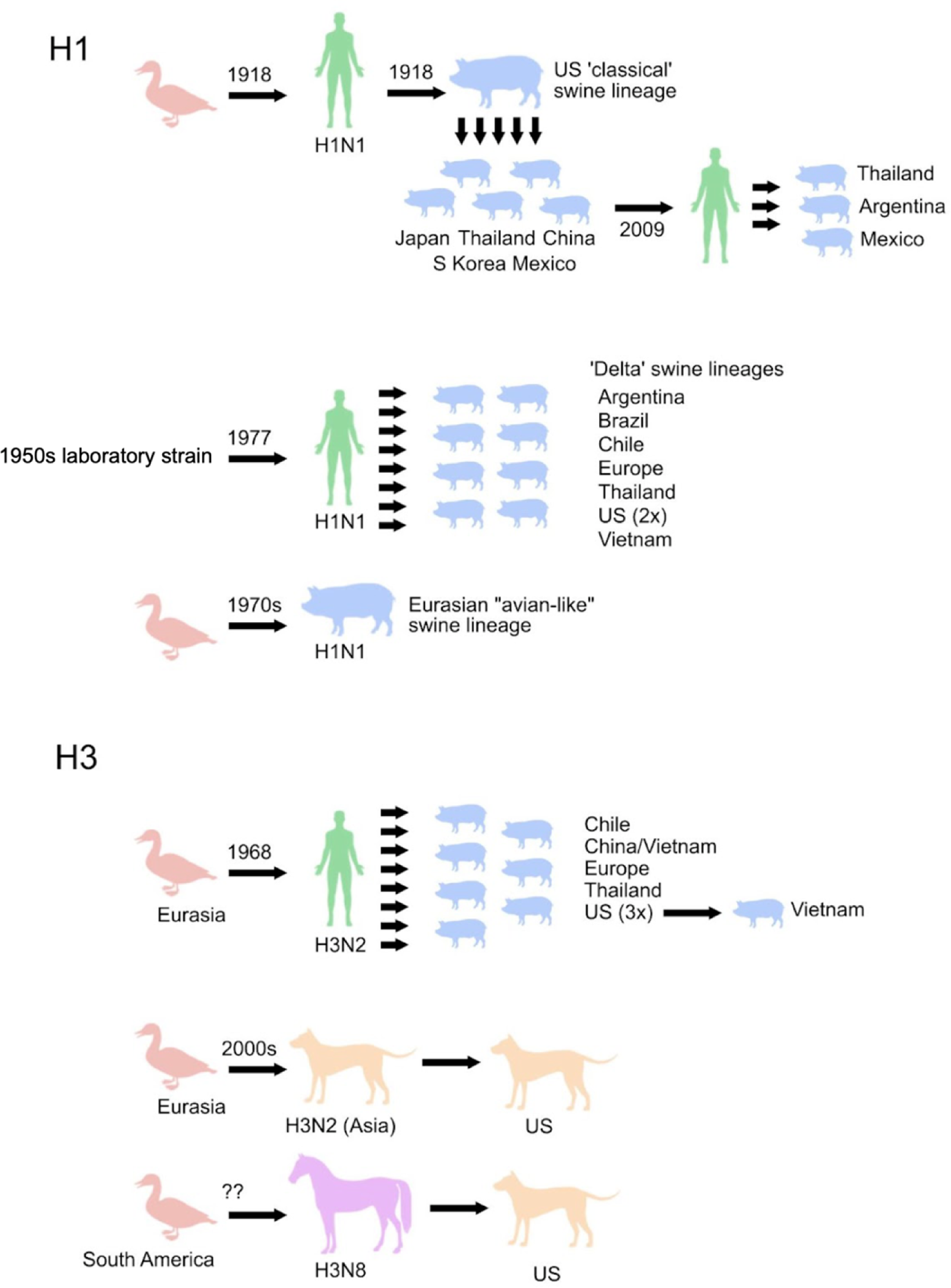
Host- and location-specific lineages of influenza H3 and H1 viruses. In 1918 influenza H1N1 viruses jumped from birds to humans causing the 1918 influenza pandemic. This viral lineage circulated in humans until 1957, and was introduced into the US swine population, giving rise to the US Classical swine lineages, which were disseminated worldwide. In 2009, due to long-distance live swine trade and reassortment events, a new lineage emerged causing the H1N1 2009 pandemic in humans, which was also reintroduced into the swine population globally. In the 1970s, the Eurasian swine lineage emerged from a direct jump from birds. In 1977, a H1N1 lineage resembling those circulating between 1946 and 1957 caused a pandemic. During its circulation, it jumped into the swine population on multiple occasions and locations, leading to the emergence of the Delta swine lineages worldwide.

Avian influenza H3 viruses are estimated to have been introduced into the equine population during the 1950s. In the 2000s, the equine lineage jumped into canines in the US. Almost simultaneously, an avian lineage was independently introduced into the dog population in Asia, and was later imported into the US. In 1968, an introduction of bird viruses into the human population caused the 1968 H3N2 pandemic, and were further passed and introduced into swine populations around the globe, with swine lineages in the US being the source of viruses circulating in pigs in Vietnam.

A combination of Bayesian mixed effects molecular clock modeling and stochastic substitution mapping was used to estimate the rates of nonsynonymous and synonymous change in several sets of positions in all lineages described above. Consistent with previous studies [29], dN/dS is higher in the head domain than the stalk domain in all hosts (Fig. 2). In both H3 and H1, the highest dN/dS in the head domain is found in the human viruses (Fig. 2, Supplementary Tables 10-13). The next highest in H3 is found in canine viruses, which unexpectedly have the highest value in the stalk. Equine viruses also have a high value for the head. The dN/dS estimates tend to be lower in swine and avian viruses of either subtype. In human viruses, dN/dS is higher in H3 than H1 in both domains.

**Fig. 2:**
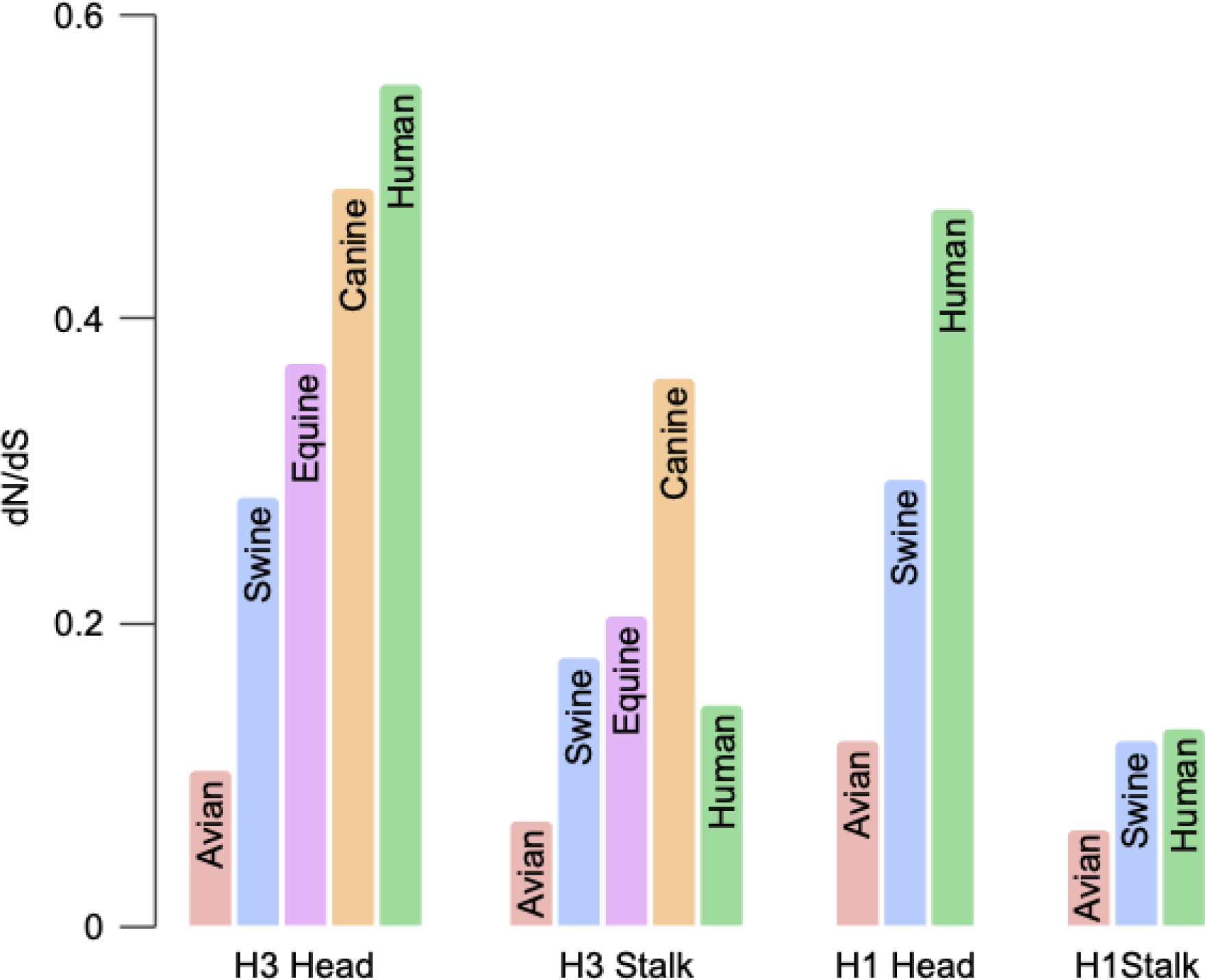
dN/dS for the head and stalk domains of H1 and H3 viruses circulating in multiple hosts.

### Synonymous Rates of H1 and H3 Across Host Species

Absolute rates of sequence change can be assessed because BEAST estimates branch lengths in units of time, which is possible because of the different isolation dates of the viruses. Higher absolute rates of synonymous change (substitutions/site/year) were observed in H3 viruses in swine, human and avian hosts, and lower in equine and canine lineages (Fig. 3). As expected, they were similar across codon subsets (Fig. S3). They were generally slightly lower in H1 than in H3, but patterns across host species were similar, with higher rates for swine and avian viruses (Fig. S4).

**Fig. 3:**
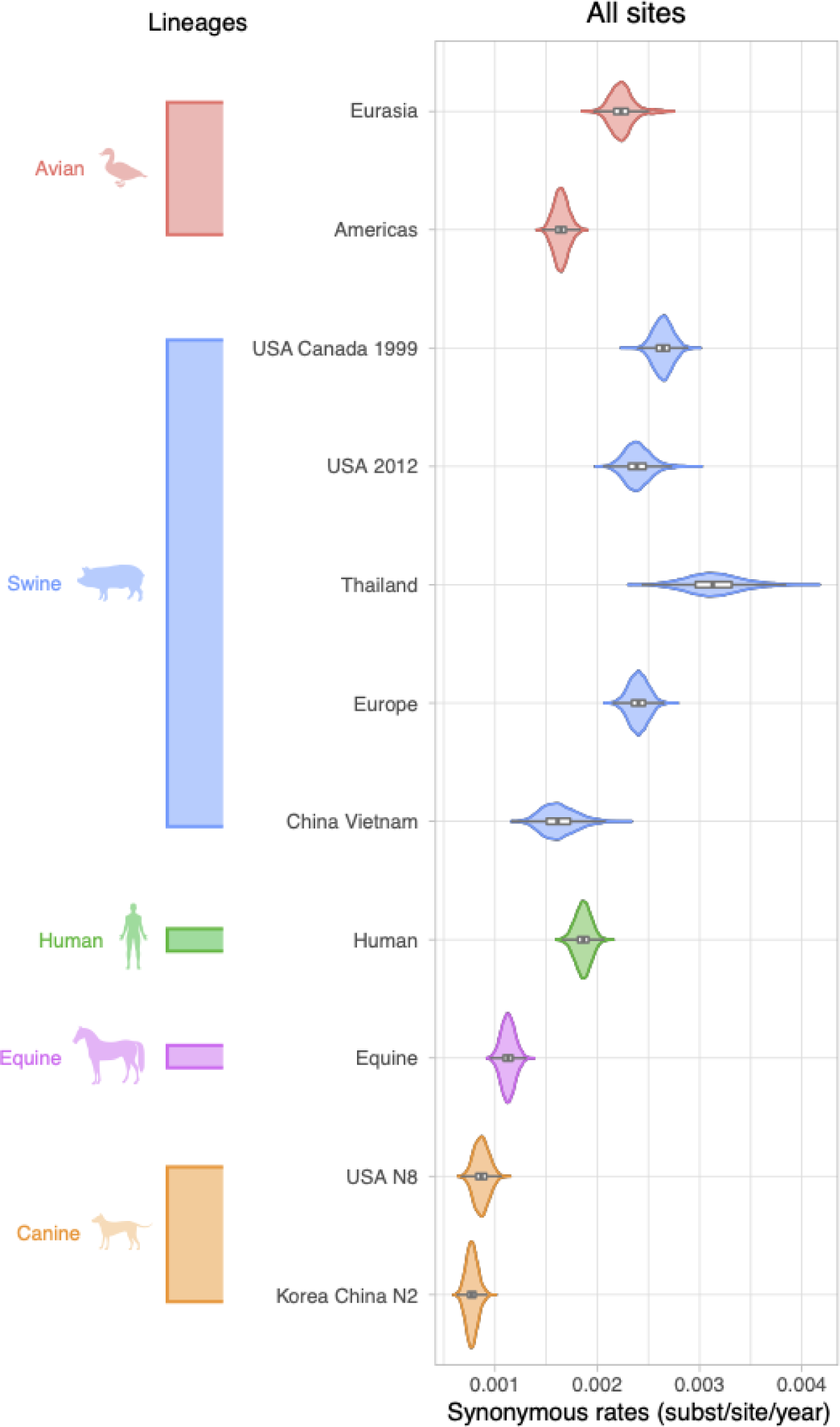
Synonymous rates for all H3 sites across lineages. Viral lineages are colored by host species: green (human), blue (swine), red (avian), purple (equine), and orange (canine). Lineage estimates with narrower violin plots (lower probability of observations taking a given value) are presented in Fig. S4 (H3) and Fig. S5 (H1).

### Protein Evolution of H3 Across Host Species

The estimates of the ratio of non-synonymous to synonymous substitutions (dN/dS (*ω)*) were higher in mammalian viruses than avian viruses across all subsets of the H3 sequence (Fig. 4). As expected, *ω* was approximately 1-4 times higher in the H3 head compared to the stalk domain (Fig. S5, and Tables S3-S6). Estimates for *ω* in antigenically relevant sites in the H3 were 2.2 to 13.5 times higher in humans compared to canines, swine, equines, and avian species (Table S4). Second to human viruses, Canine Asia H3N2 and Canine USA H3N8 lineages had the highest *ω* estimates across all sequence subsets. However, we observed a strikingly different pattern in the stalk domain. Estimates for *ω* in the stalk domain are 1.8 to 7.4 times higher in canines Table S6) compared to human, equine, swine and avian lineages. We explore this observation in more detail below.

**Fig. 4:**
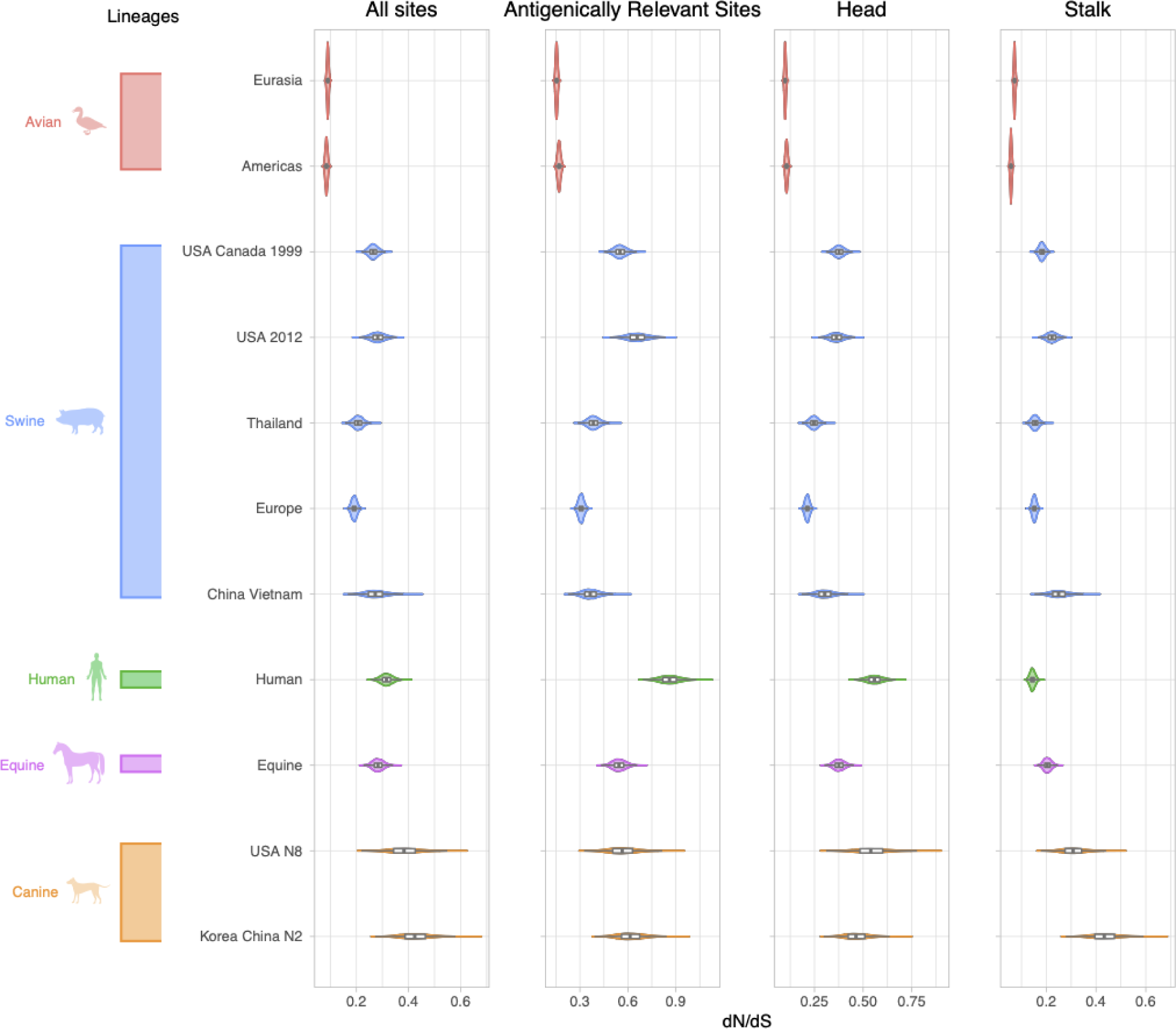
dN/dS for all H3 sites, antigenically relevant sites, head and stalk domains across lineages. Viral lineages are colored by host species: green (human), blue (swine), red (avian), purple (equine), and orange (canine). Lineage estimates with narrower violin plots (lower probability of observations taking a given value) are presented in Fig. S5.

### Protein Evolution of H1 Across Host Species

Patterns of *ω* in H1 viruses were generally similar to those observed for H3. Again, the estimates of *ω* were higher in H1 mammalian viruses than in H1 avian viruses in all subsets of positions (Fig. 5). As expected, *ω* was approximately 1.5-5 times higher in the H1 head compared to the stalk domain (Fig. S6, and Tables S7-S9). We estimated the highest overall *ω* for the H1 human lineage that circulated from 1918-1957 for all sites and head domain (Fig. 5, Tables S7 and S8).

**Fig. 5:**
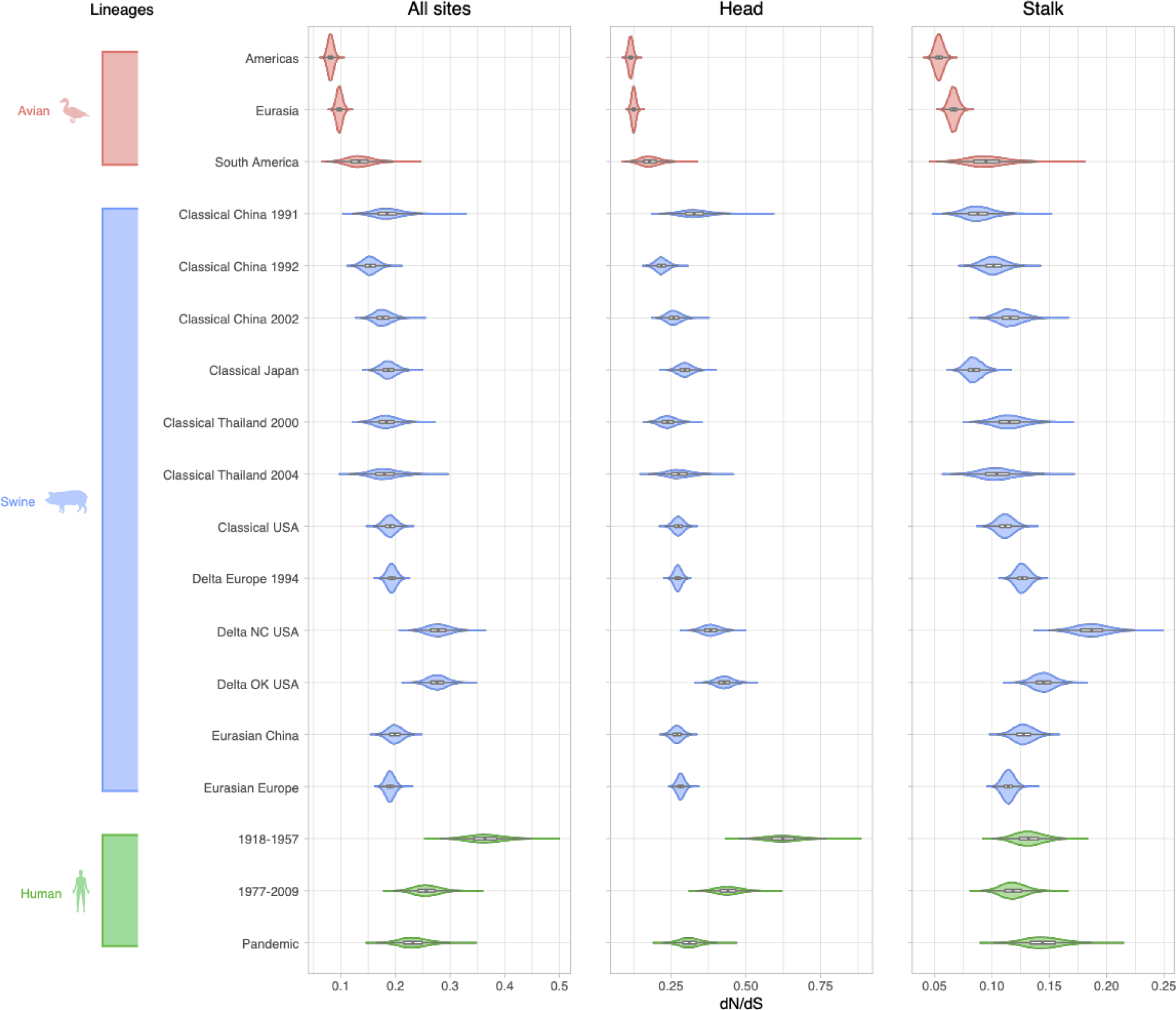
dN/dS for all H1 sites, head and stalk domains across lineages. Viral lineages are colored by host species: green (human), blue (swine), red (avian). Lineage estimates with narrower violin plots (lower probability of observations taking a given value) are presented in Fig. S6.

Estimates for *ω* for H1 head domain were 1.5 to 5.8 times higher in humans than in swine and avian species (Table S8).

The *ω* of the head domain of human-like “delta” swine lineages circulating in the USA in 2005- 2019 was between 1.4 to 1.9 times higher than several other swine H1 lineages, including delta, classical, and eurasian swine viruses (Fig. 5 and Table S8). The US delta swine viruses also had an elevated *ω* in the stalk domain compared to other lineages (Fig. 5 and Table S9).

### Comparison of Nonsynonymous and Antigenic rates of evolution

Rates of antigenic evolution (antigenic units per unit time) of human and swine HA, derived from hemagglutination inhibition (HI) assays [47, 48], are compared to nonsynonymous rates per unit time in Fig. S7. In all cases in which precision permits comparison, human H3 and H1 have higher antigenic rates than their swine counterparts. Human H3 has a higher nonsynonymous rate per unit time than swine H3, but human H1 has a lower rate than many types of swine H1. The large uncertainties of antigenic rate estimates for some types of swine HA limit the value of comparisons.

### Glycosylation Patterns Across H3 Lineages

We investigated trends in glycosylation for each lineage using the motif Asn-X-Ser/Thr-X, where X is any amino acid other than proline. The average yearly number of glycosylations for H3 (Fig. 6) varies markedly across host species with an average of 6.2 glycosylations in avian lineages, 7.0 in canine, 7.8 in equine, 9.2 in swine, and 11.5 in humans (File S3). While the number of glycosylations in avian, canine, and equine hosts generally remained constant over time, the number of glycosylations in human and swine viruses increased over time. The human H3N2 lineage inherited 7 glycosylations from its avian ancestor. It subsequently gained 6 glycosylations (Table 1), but the avian lineage did not gain or lose glycosylations in the decades that followed. During 1968-2019, the human H3 lineage gained an average of 1.1 glycosylation per decade, peaking at 13 glycosylations in 2017.

**Fig. 6:**
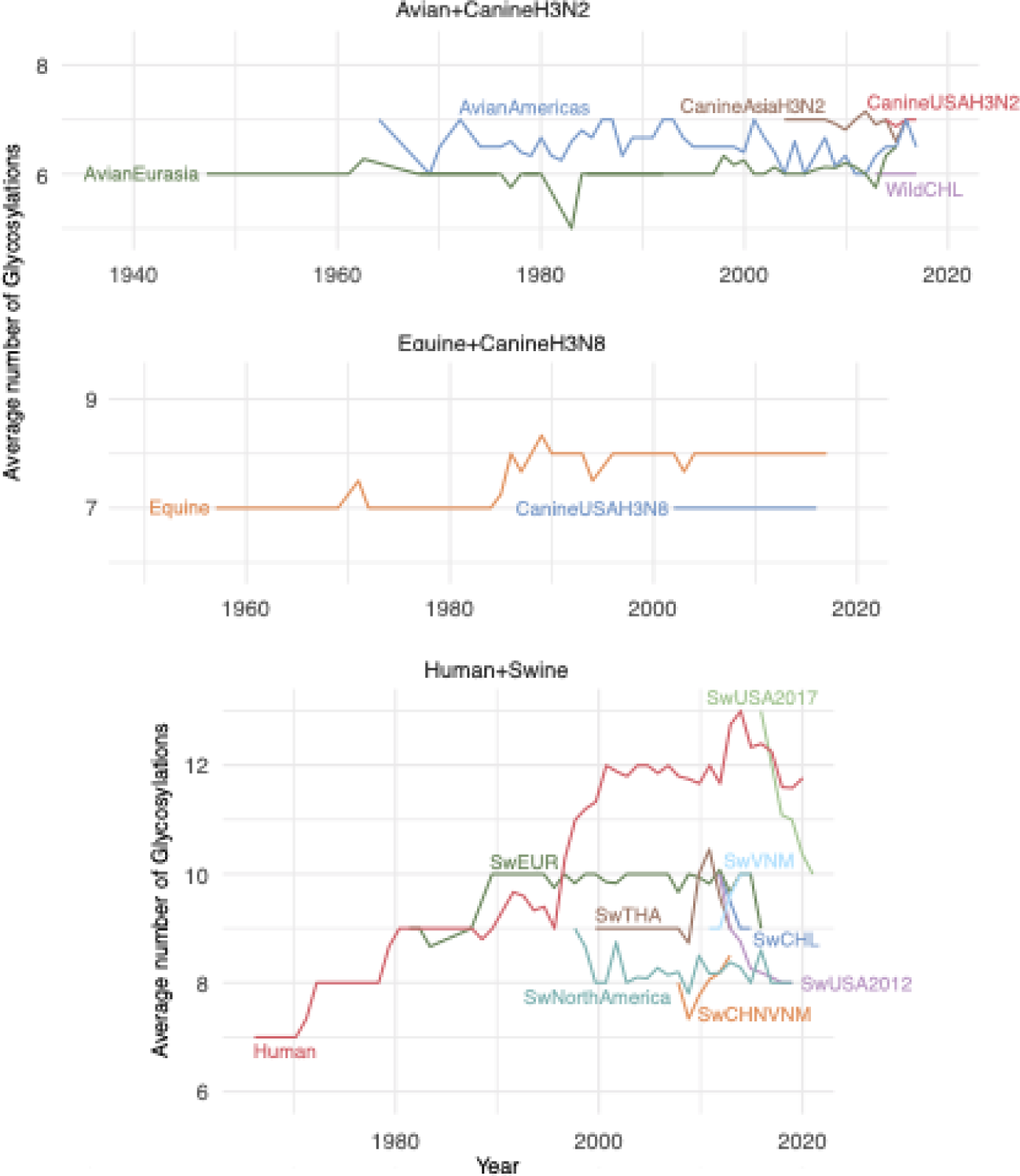
Average number of H3 glycosylations over time. For each lineage the average number of glycosylation sites is plotted for each calendar year for which numbers are available. The average is taken over all isolates and internal nodes (reconstructed ancestral sequences) dated to that year.

**Table 1.** Average number and rate of accumulation of glycosylation in H3 lineages

| **Lineage** | **Slope of the annual average number of glycosylations** | **Annual average number of glycosylations** |
| --- | --- | --- |
| Swine Chile | -0.90 | 9.17 |
| Swine USA 2017 | -0.66 | 11.26 |
| Swine USA 2012 | -0.16 | 8.30 |
| SwineNorAm | -0.05 | 8.23 |
| Avian Americas | 0.00 | 6.47 |
| Avian Chile | 0.00 | 6.00 |
| Canine Asia H3N2 | 0.00 | 6.96 |
| Canine USA H3N2 | 0.00 | 6.95 |
| Canine USA H3N8 | 0.00 | 7.00 |
| Swine Europe | 0.00 | 9.89 |
| SwineThailand | 0.00 | 9.35 |
| Avian Eurasia | 0.01 | 6.08 |
| Equine | 0.02 | 7.76 |
| Human | 0.11 | 11.54 |
| Swine China, Vietnam | 0.19* | 8.01 |
| Swine Vietnam | 0.21* | 9.58 |

\* Only 3-4 years of data is available for these lineages.

Subsequent host-switches of H3N2 viruses from humans to swine seeded additional human-like H3 lineages in pigs. After establishment in swine, the number of glycosylations typically decreased over time to between 8 and 9 glycosylations (Fig. 6 and Table 1). Two swine lineages in China and Vietnam gained glycosylations over time, but there are only a few years of data available for these lineages and long-term trends are unclear.

The avian H3 lineages have stably maintained the lowest number of glycosylations (6 to 7) in both the Americas and Eurasia. The equine lineage maintained a low glycosylation average of 7 for approximately 30 years after emergence, but in the late 1980s it gained a single glycosylation that has been maintained as of 2019 (Fig. 6). One site was lost in the genesis of canine H3N8, which mostly maintained the resulting glycosylation state. The avian-origin canine H3N2 clade has also maintained 7 glycosylations, like the Eurasian avian lineage from which it derived (Fig. 6), though one of the glycosylations is at a different position.

In H3, glycosylations are accumulated in the HA structural domains at different rates across lineages (Figs. S8 and S9, and Table S14 and S15). Glycosylations were mainly gained and lost in the HA stalk domain in Avian viruses circulating in the Americas (Fig. S9), with no net change in the number (Table S15). There were gains of 1-2 glycosylations in the stalk domain of swine viruses circulating in China and Vietnam. The number of glycosylations in the HA head domain of human viruses increased by 3 during the period between late-1990s and early-2000s, and was overall stable during approximately 15 years, followed by the loss of at least one glycosylation. The stalk domain of the human lineage kept a stable number of 5 glycosylations from its emergence to 2010, but since then has gained 2 glycosylations.

### Glycosylation Patterns Across H1 Lineages

The average number of glycosylations (Fig. 7, Table S16 and File S4) varies across H1 lineages in a manner similar to H3. The avian lineages have maintained a stable, lower number of 7 glycosylations over time (Fig. 7). Like human H3, the human 1918-1957 H1 lineage, which is thought to have originated in birds, initially had 7 glycosylations. By 1957, the lineage had 11 glycosylations, a net gain of 4 (Fig. 7 and Table S16). As expected, the human 1977-2009 H1 lineage, which began with a virus genetically similar to H1N1 from the 1950s, inherited 11 glycosylations. This number dropped to 10 by the time the virus went extinct in 2009 during the swine-origin H1N1 pandemic. The currently circulating swine-origin H1N1 pandemic (PDM) lineage in humans inherited a relatively low number of glycosylations from swine (n = 8), but this number increased to 9 within a decade (Fig. 7 and Table S16). Surprisingly, the initial classical swine virus circulating in North America experienced a dramatic gain in glycosylations since 1970. The virus emerged in US swine during the 1918 pandemic with 7 glycosylations, like early human H1N1, but this number increased to 11 by 2019, reaching the same maximum as humans. In contrast, the Eurasian avian-like H1 swine lineage only gained 1 glycosylation in the decades since the virus switched hosts from birds to pigs in the late 1970s. The swine delta lineages (Fig. 7) resulted from host jumps from Human1977-2009 viruses. The average number of glycosylations in swine delta lineages and PDM lineages has remained relatively stable over time (Table S16). There have been multiple spillovers of human pandemic H1N1 into swine since 2009.

**Fig. 7:**
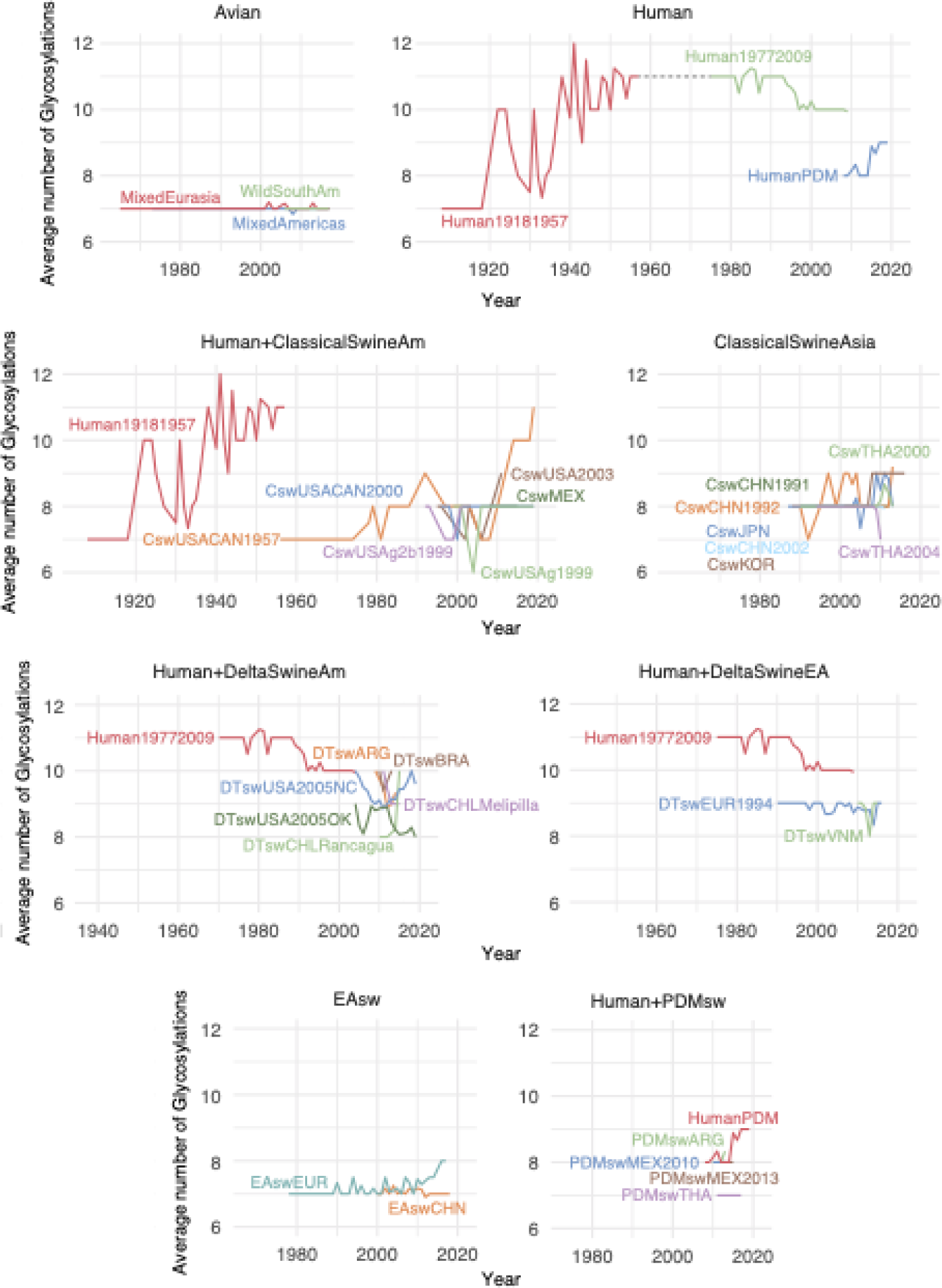
Average number of H1 glycosylations over time. See Fig. 6 legend for details.

Between 1992 and 2005, six of the swine lineages (DTswEUR1994, DTswVNM, DTswUSA2005OK, DTswCHLRancagua, DTswCHLMelipilla, EAswEUR) have mostly lost glycosylation at position 94 (File S4).

As in H3, the accumulation of glycosylations in H1 occurred more frequently in the head domain in the human lineages (1.5-5.8 times more for H1 human lineages than in other swine and avian lineages) (Figs. S10 and S11, and Tables S17 and S18), whereas swine tended to gain and lose glycosylation at similar proportions in either HA domain, irrespective of subtype.

No gains or losses in glycosylation were observed in the stalk domain of human H1 lineages, avian, and pandemic swine viruses (Fig. S11).

### Evolution of the H3N2 and H3N8 Canine Lineages

Canine viruses form two clades in the H3 phylogeny. One consists of H3N8 viruses and was introduced into canines from equines in the 2000s in the United States (canine USA H3N8 lineage). The second consists of H3N2 viruses and was introduced into canines from Eurasian birds in the 2000s in Asia (canine Asia H3N2 lineage) and subsequently disseminated to the US around 2015, establishing the Canine USA H3N2 lineage. The existence of two independently derived canine clades allows confirmation that some evolutionary patterns are related to canine hosts, rather than being idiosyncratic features of a sequence type.

A total of 14 amino acid changes were inferred to occur on the branches containing the host-switches to canines from avians or equines (Fig. 8).

**Fig. 8:**
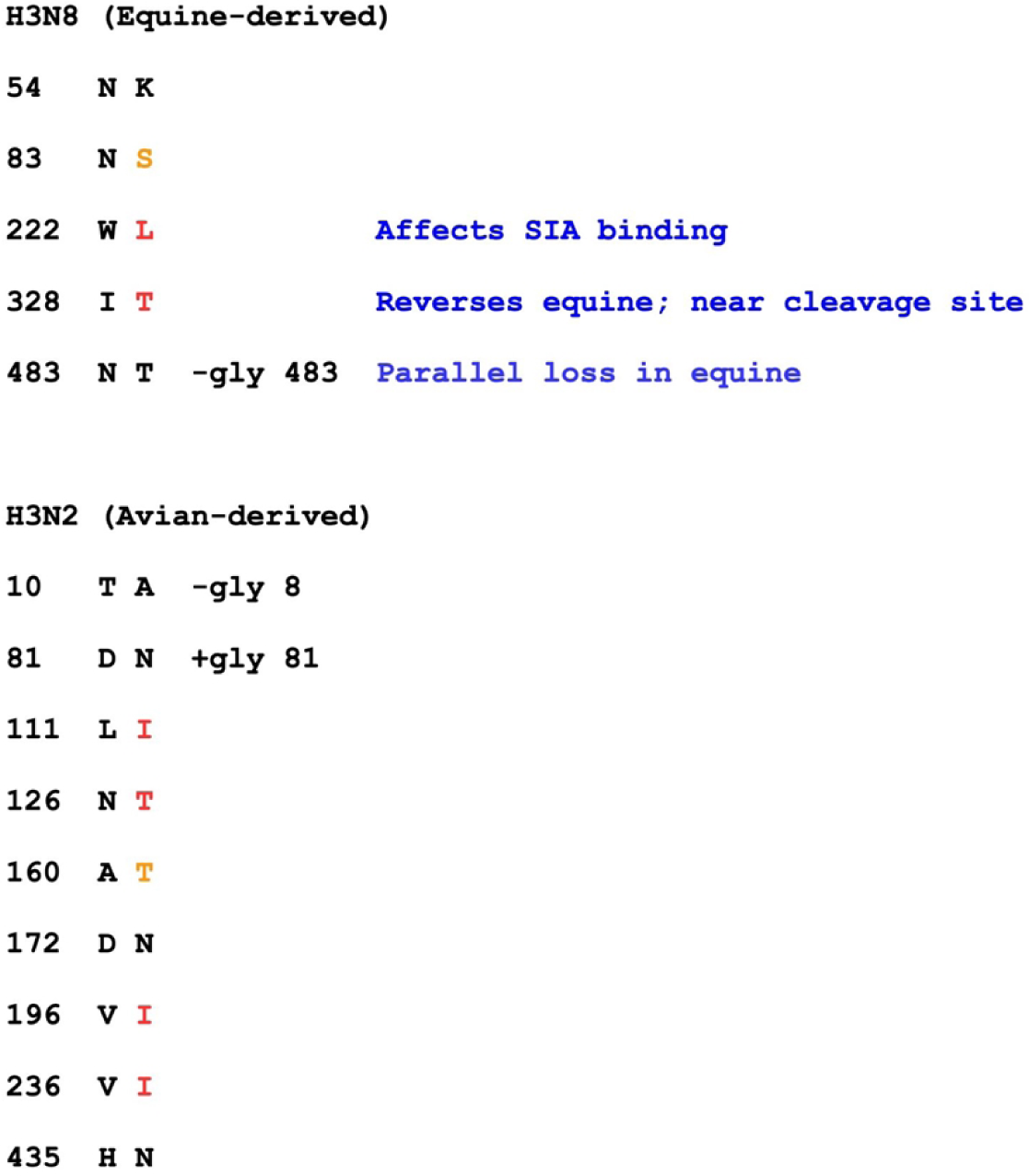
Amino acid changes on host-shift branches leading to canine clades

On the host-switch branch leading to the canine USA H3N8 lineage, there are five nonsynonymous changes and one synonymous change. The 5:1 nonsynonymous:synonymous (N:S) ratio on this branch is statistically distinguishable from the ratio in the ancestral equine lineage (366:537 = 0.68:1; p=0.044, Fisher’s exact test), but not from the ratio in the derived canine clade (101:111 = 0.91:1; p=0.11). Some nonsynonymous changes occurred in sites with known functional importance. One of the substitutions (W222L) is located adjacent to the sialic acid binding site. I328T is a reversal of an equine-specific change: the equine sequences have mostly I and occasionally L (6%), but 97% of other H3 viruses have T, and none has I or L. This position immediately precedes Arg329, after which proteolytic cleavage splits HA into HA1 and HA2 (the peptide bond between 329 and 330). The N483T change eliminates a glycosylation site that was present in the equine lineage (confirmed by crystal structures, PDB entries 4uo0 and 4unw) (Fig. 8). Independent loss of this site occurred in a close equine relative of the ancestor of the CanineUSA H3N8 virus within about three years (according to the MCC tree, 95% HPD branch lengths) and possibly simultaneously (Fig. S12). The parallel losses are clearly distinguishable: that on the host-shift branch results from a second-position change of codon 483 from AAT (asparagine) to ACT (threonine), while the other results from a third-position change to AAG (lysine). This conclusion comes from the maximum parsimony reconstruction, but is also supported by the sequences of the earliest CanineUSAH3N8 isolate and the affected equine isolate, which bear ACT and AAG, respectively.

The canine Asia H3N2 host-switch branch contains 9 nonsynonymous and 19 synonymous changes. The 0.47:1 N:S ratio is intermediate between that in the progenitor avian Eurasia lineage (472:2217 = 0.21:1) and that in the derived canine H3N2 clade (147:157 = 0.94).

However, it is not statistically distinguishable from either due to the low absolute counts. The canine clade and the avian progenitor are clearly distinguishable from each other (p=2e-30, Fisher’s exact test).

No changes at the same amino acid positions occur on the two host-shift branches, but in some cases (indicated by red in Fig. 8) the amino acid resulting from a change in one is identical to that inherited at the same position by the other canine clade. In two additional cases (indicated by orange), a change results in a serine at a position where the other last common ancestor (LCA) has the chemically similar threonine, or vice versa. In both cases the observed change is due to a transition mutation, whereas a change to the amino acid present in the other LCA (threonine or serine, respectively) would require a transversion of a type that is much less frequent, as determined by analysis of fourfold-degenerate synonymous sites.

A total of 21.4% of the amino acid changes on these branches result in changes of glycosylation state. This is larger than the fraction (157/3755, or 4.1%) for the remainder of the tree (p=0.019, Fisher’s exact test). A similar conclusion holds for the ratio of glycosylation changes to synonymous changes (p=0.0070, Fisher’s exact test).

The non-canine ancestors of both canine clades bear two glycosylation sites near the membrane-proximal portion of HA, at positions 8 and 483 (HA2 154). One of these sites is lost in the genesis of the canine H3N2 clade (8), and the other is lost in the genesis of canine USA H3N8 (483). Canine USA H3N8 inherits a glycosylation site at position 63, which is absent from H3N2, and the canine H3N2 clade gains a glycosylation at 81. These sites lie at nearly the same position along the HA axis, and may cover similar regions of the protein’s surface (Fig. S13).

Additional evidence for their interchangeability comes from human H3N2, in which a gain of glycosylation at 63 is quickly followed by a loss of glycosylation at 81 in the “trunk” of the tree.

### Changes within Canine Clades

Fig. 9 shows the distribution of amino acid changes within the canine clades across the protein sequence. The variance of these counts is too large to be explained, except by extremely improbable occurrences, unless the rate at some of the sites exceeds the neutral expectation substantially. It would be explained if the counts for the seven sites with at least four changes reflected high rates, implying positive selection, rather than chance events. Unless rates at the remaining sites were mostly close to the neutral expectation or to zero, more sites would have to be subject to positive selection. Positive selection on a site might be obscured by purifying or weak selection in other parts of lineages, as appears to be the case for these changes (see below).

**Fig. 9:**
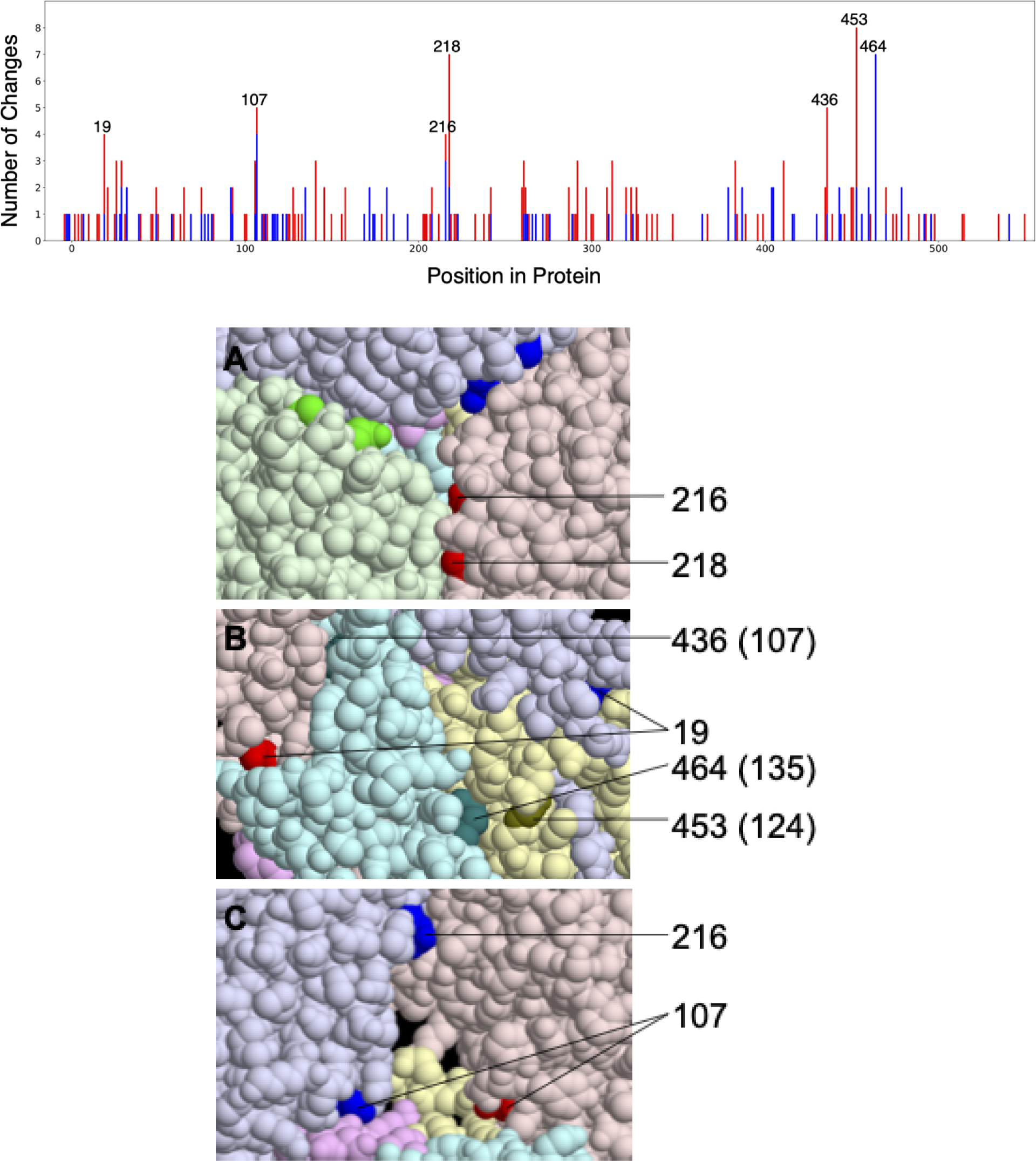
Amino acid changes across the H3 protein of canine isolates. Total number of amino acid changes across the H3 protein of canine isolates (top). Changes in Canine H3N2 (red) and Canine USA H3N8 (blue) are stacked. The seven positions with at least four changes are labeled. Six of these are interface positions in all monomers of all four structures considered (PDB entries 4uo4, 6n4f, 4uo0, and 4unw). The seventh (464) is an interface position in all monomers of the two equine structures. Interface positions can be found in File S5. The seven positions evolving most rapidly in canine H3 shown on the HA structure (bottom). Three views together show all seven positions. The six protein chains (three each of HA1 and HA2) are given different light colors, and the seven positions are shown in darker versions of the same color. In panel A, three 216 and 218 residues appear (one of each on each monomer), but only one of each is labeled. In panel C, one of the HA monomers (chain C) is made invisible to reveal the buried position 107. Positions 216 and 218 in chain A are not colored in this view because they are at the interface with the invisible chain C. HA2 positions are given in parentheses. The HA structure is that of A/Equine/Richmond/07, PDB entry 4UO0.

The seven most rapidly evolving positions, those with four or more total changes in the canine clades, are 19, 107, 216, 218, 436 (HA2 107), 453 (HA2 124), and 464 (HA2 135) in the conventional H3 numbering (based on human H3 after removal of the 16-residue leader peptide). We refer to this set of positions as C7.

The locations of these seven positions in hemagglutinin crystal structures appear to lie disproportionately at interfaces between protein chains. This impression is confirmed by a quantitative analysis. We define interface residues as those for which any atom is within 5 angstroms of an atom of a different protein chain. This includes the interface between HA1 and HA2 chains derived from the same full-length HA. The set of positions meeting this criterion varies among HA crystal structures, presumably in part because of differences in the protein sequences, and also between monomers not related by crystallographic symmetry. We consider four structures: one from each canine clade (PDB entries 4uo4 for CanineUSAH3N8 and 6n4f for CanineAsia/USAH3N2), and two of equine hemagglutinins (4uo0 and 4unw), which retain some ancestral residues that have changed in the CanineUSAH3N8 viruses.

Of the seven rapidly changing positions, six (all but 464) are interface positions in all monomers of all four structures. All seven are at interfaces in at least one monomer of these structures. The corresponding fractions for all positions in the protein are ∼35.5% (interface in all monomers of all structures) and ∼42.9% (interface in at least one). Thus, the probability that at least six of seven positions chosen at random lie at interfaces in all structures is 1.0%, and the probability that all seven do in at least some cases is less than 0.3%.

This result is not due to more rapid evolution of interface residues in general. In fact, in non-canine lineages, interface positions evolve more slowly on average than non-interface positions by nearly a factor of two. The C7 positions are also not particularly rapidly-evolving in other lineages. Thus, the over-representation of interface residues among the rapidly-evolving sites is even more striking than suggested by the probabilities given above.

This phenomenon is not restricted to the seven most rapidly-evolving sites. At the remaining interface positions, the average number of changes (0.44) is larger than that at non-interface positions (0.37) (p=0.041, randomization test). In non-canine lineages, the reverse is true. The ratio of interface changes to non-interface changes, excluding the C7 sites, is higher in canines by a factor of 2.6 (p=1e-10, 95% CI=(1.9, 3.5), Fisher’s exact test).

The structures demonstrate that changes at position 135 of HA2 (464 of HA), the only C7 position that is non-interface in some of the structures, can have large effects on inter-subunit interfaces. Changes at this position in canines are restricted to the equine-derived H3N8 clade. Both equine structures have the ancestral glycine at this position. It lies within 5 angstroms of the associated HA1, and within 5.5 angstroms of the Arg at 124 (another C7 position) of another HA2 chain. The CanineUSAH3N8 structure, in contrast, has an aspartate at position 135, and it is further than 5 angstroms from any other chain. The backbone configuration surrounding this position is altered, and the positioning of several nearby residues is affected. Likely as a consequence, an inter-chain bidentate salt bridge between Glu131 and Arg163, found in both equine structures, does not form in the canine structure.

Changes of Arg124 will also eliminate this salt bridge. The ancestral glycine at position 218 has backbone torsion angles, corresponding to right-handed beta-strand, that are “forbidden” for non-glycine residues, which suggests that changes there will disrupt the structure. At five of the seven positions, the ancestral residue is inferred to have changed to several different residue types, which is more consistent with disruption of favorable interactions than creation of new ones.

## DISCUSSION

Studying the evolution of influenza sequences in non-human hosts is important for several reasons: the burden of non-human infections, the ever-present possibility of a jump to human hosts, the knowledge gained about viral evolution in general, and the light it sheds on evolution of human influenza. Due to the paucity of antigenic data for non-human influenza, a molecular evolutionary approach is particularly important.

To study the evolution of H1 and H3 in all major hosts, we first built phylogenetic trees for representative sets of coding sequences. We determined rates of nonsynonymous and synonymous change in the sequences, and in portions corresponding to the two domains of HA. We also reconstructed the specific changes that occurred on all branches of the tree.

The ratio of nonsynonymous to synonymous changes, relative to the ratio expected in the absence of selection, reflects the selective forces acting on the protein sequence. Fig. 2 shows this quantity for all hosts for the head and stalk of H1 and H3, and Tables S10-S13 compare them among hosts. In all cases, the avian dN/dS is lower than all of the mammalian values.

Where the mammalian values are statistically distinguishable from each other, the human value is highest, except for the special case of canine stalk. Where differences exist, they might reflect differences in the strength of purifying selection, differences in positive selection, or some combination of the two.

High dN/dS on the head of human HA has long been attributed to selection for antigenic change, which allows escape from antibodies elicited by previous infections. Values for the head in other mammalian hosts are indistinguishable from human dN/dS or intermediate between human and avian values. The simplest explanation of this pattern is that there is selection for antigenic change in these other mammals as well, albeit weaker, but little or none in avians.

Influenza infection in birds differs from that in mammals, mainly affecting the gastrointestinal tract, and birds may be less likely to be reinfected with the same subtype in their lifetimes. These differences, and other possible explanations of weak antigenic selection, may be due to the long association of these subtypes with avian hosts [5].

The N:S ratio on the H1 and H3 stalks is also higher in mammalian than avian viruses. The very high canine value can be explained by adaptations to canines, discussed further below. The higher value in other mammals might reflect weaker purifying selection or the operation of positive selection. There is no obvious reason for purifying selection on the stalk to be generally weaker in mammals, but the possibility cannot be excluded. Adaptive evolution of the stalk might involve escape from rare anti-stalk antibodies. The same force responsible for higher mammalian rates in the head would then also explain higher rates in the stalk. This possibility would be a concern for universal influenza vaccines designed to elicit antibodies that target the HA stalk [49].

Some significant differences exist in N:S ratios among lineages of the same subtype that infect the same host (Tables S5, S6, S8, and S9). The dN/dS of the head is lower in the 2009 human H1N1 pandemic lineage than in the other human H1N1 lineages. This likely reflects the small amount of time since the introduction of the 2009 lineage into humans and the consequent weakness of selection for antigenic change over much of its history. The vaccine strain for 2009pdm was not updated until 2016 because of a paucity of antigenic change. There is significant variation among both H3 and H1 swine lineages on both the head and the stalk. Differences in agricultural practices and swine vaccination rates over time and among countries may explain much of this variation.

We analyzed changes at 131 H3 positions commonly described as “antigenically relevant” to allow comparison to other results. These positions are based on dubious premises: that all changes in the membrane-distal portion of the HA head that became widespread among human H3N2 were caused by selection for antigenic change, and that regions of the protein not observed to have changed are not bound by antibodies. Analysis based on these sites can lead to circular reasoning, since positions that have changed in one part of the tree are more likely to change in other parts, even if the changes are not due to antigenic selection. This potential for this effect is highest in the human lineage, since these sites were selected based on human virus data.

Estimates of the rate of antigenic change, based on laboratory measurements, are available for human and swine H1 and H3 (Bedford et al. 2014; Lewis et al. 2016). Large uncertainties for several of the swine rates limit the conclusions that can be drawn from comparisons (Supplementary Fig. 7). In all cases where a conclusion is possible, the rate of change in human viruses is larger than that in swine. This is consistent with the generally higher rate of nonsynonymous change on the head, relative to the synonymous rate, in human viruses, and its interpretation as an indication of stronger selection for antigenic change. Perhaps more appropriate is consideration of the rate of nonsynonymous change per unit time. By this measure, human H3 again has a higher rate than swine H3, but human H1 is slower than most types of swine H1. The implication is that there is more antigenic change, on average, for a nonsynonymous change in human than swine H1. This is expected if there is stronger selection for antigenic change. Even in the absence of such selection, HA sequences, like those of most proteins, are expected to change over time. This will result in some antigenic change, but less than that due to amino acid changes selected for the antigenic difference that they cause.

The number and locations of glycosylation sites vary across the HA trees, especially on the head domain. Glycosylation of the head can shield the protein from antibodies [50–52]. It also comes with various costs [31,52,53]. The optimal glycosylation state reflects a trade-off between the costs and the benefits, both of which may vary considerably among hosts and due to other factors.

As previously described [30,52,54], there has been net acquisition of glycosylations on the head since the introductions of H1 and H3 to humans in 1918 and 1968. This phenomenon is attributed to selection for the shielding effects of glycosylation. The 2009 pandemic H1N1 may be in the early stages of a similar trajectory, having gained a head glycosylation after a few years. In contrast, avian H1 and H3 have generally maintained just one glycosylation on the head over long periods of time.

Equine and Canine H3 typically have two or three head glycosylations. Swine H3 lineages converge to around four, despite being derived from human progenitors with a variable number. Loss of glycosylation is common in the swine lineages derived from highly-glycosylated human H3 and on the host-switch branches leading to them. Presumably this reflects weaker antigenic pressure in swine compared to humans, such that the costs of glycosylation outweigh its benefits, leading to selection for loss.

The pattern in swine H1 is similar in that the average number of head glycosylations is intermediate between the low number in avians and the higher numbers attained over time in humans. There is, however, considerable variation among swine lineages, mostly ranging from a single head glycosylation (like avian viruses) to three. The head has no glycosylations for a few years in the avian-derived EAswEUR. As discussed above, N:S ratio also varies significantly among swine lineages.

The overall pattern of head glycosylation is similar to that of nonsynonymous to synonymous rate on the head: highest in human viruses, lowest in avian, and intermediate in non-human mammals. Differences in antigenic pressure among hosts would explain both patterns.

Considering only the total number of glycosylations hides the complexity of their evolution. This complexity is exemplified by position 94 of H1. Loss of this glycosylation site occurs in the ancestors of most isolates of six swine lineages. The site is present throughout most of the tree, including the avian lineages, so it would seem to have a cost that is specific to swine. Nevertheless, it is regained six times in EAswEUR, suggesting that it has a net advantage in swine under some circumstances. It is never regained, however, in EAswCHN, which descends from EAswEUR. The explanation may be that glycosylation at position 94 provides protection against antibodies in those swine that have received certain vaccines [45, 55]. Vaccination of swine is not common in China [56].

There were two independent introductions of H3 to canines. An H3N8 clade derives from equine H3N8, and an H3N2 clade derives from avian H3N2. There is evidence of adaptation of the HA sequence to canines within these clades and on the branches leading to them.

There are five nonsynonymous changes and one synonymous change on the branch containing the jump from equines to canines. This 5:1 ratio is high compared to the neutral expectation of ∼2.38:1, and higher still in comparison to the ratio on the remainder of the tree. This might suggest positive selection. However, the absolute numbers are so low that there is much uncertainty in the ratio. It is statistically distinguishable from the ratio in the ancestral equine lineage, but not from that in the derived canine clade. The host-switch branch for H3N2 canine is longer, which might obscure any burst of nonsynonymous changes associated with the change of host. Nevertheless, the nature of the changes on these branches suggests that some of them are adaptations to canines.

There are three changes of glycosylation state on these branches. The fraction of the total amino acid changes that change glycosylation state is higher on these branches than in the remainder of the tree, as is the ratio of glycosylation changes to synonymous changes.

Furthermore, these changes are convergent in two senses. First, the resulting canine HAs carry the same number of glycosylation sites, whereas the numbers on their parents differ by one.

Second, though the changes result in two sites present in each canine LCA and absent from the other, in each case there is a membrane-proximal site and a more distal site at approximately the same distance from the membrane (Fig. S13). These changes are likely adaptations to canine hosts.

Glycosylation at position 483 is lost on the H3N8 host-shift branch. This position is glycosylated almost universally throughout the H3 tree. Loss of this site occurred only three other times, each affecting a single isolate. One of these losses occurred in a close equine relative of the equine ancestor of canine N8. This pattern suggests that there was, for a time, selection for loss of glycosylation at position 483 in equine hosts, and that this loss made the equine virus better suited to canine hosts, facilitating the host jump.

Several other changes on the canine host-shift branches produce the same amino acid in one clade that is inherited by the other, or lead to a serine in one and a threonine in the other, suggesting that they are adaptations to canines. The nature of some of these changes suggests that they are functional. The W222L substitution, which occurred on the H3N8 host-switch branch, has been shown to alter the specificity of sialic acid binding [57]. The H3N2 canine LCA also has L at this position, inherited from its avian progenitor. I328T is a reversal of an equine-specific change: the equine sequences have mostly I and occasionally L (6%), but 97% of other H3 viruses have T, and none has I or L. Position 328 is close to the site of proteolytic cleavage of HA into HA1 and HA2 (just after Arg329). The presence of I or L in equine H3, and reversion to T in the formation of canine H3N8, may reflect differences in the specificities of proteases performing the cleavage in different hosts.

The nonsynonymous:synonymous ratio on the H3 stalk is much higher in canines than in any other host. The ratio on the head is higher in canines than in any other non-human host, and close to the human ratio.

Amino acid changes within the canine clades occur disproportionately at positions near interfaces between HA protein chains. This phenomenon is strongest in the seven positions with the most changes in canines (the C7 positions), but is also seen in the remaining positions. A reasonable explanation is that such changes are selected because of their effects on the rearrangement of the HA structure that occurs under the acidic conditions of the endosome and is necessary for viral entry into the cytoplasm [58]. These amino acid changes may alter the pH at which this transition occurs, bringing it closer to the optimum for canine hosts.

Natural variation and laboratory mutants alter the HA transition pH, and sequence changes of this type are thought to be involved in host adaptation [58–62]. The optimal transition pH reflects a trade-off between efficient rearrangement in the endosome and avoidance of rearrangement prior to phagocytosis. Conditions within and outside the endosome may vary among hosts, so that the optimum in a new host differs from that in the ancestral host, leading to selection for change of transition pH. Selection of this sort has been observed in laboratory adaptation of avian influenza to mammalian hosts [58].

A T30S change in the equine virus, which is inherited by canine H3N8, raises the transition pH [22]. The authors suggested that this predisposed the affected equine clade to jump to canines. The transition pH remains lower than typical for HA after this change—5.2 as opposed to ∼5.7—so selection to increase it might be expected in the derived canine clade. Based on structural information, changes at three of the C7 positions would be expected to weaken associations and hence raise the transition pH, while it is difficult to predict anything about the other four. Five of these seven positions change to more than one amino acid within canines, which is more consistent with disruption of existing interactions than production of new ones.

Nevertheless, selection for stronger subunit association might operate after a change that brings the transition pH below its optimum. In support of this notion, there are three reversals (a derived amino acid gives rise to the ancestral residue) at C7 positions.

The sequence datasets compiled in this study were subjected to substantial downsampling to mitigate the computational burden of Bayesian phylogenetic inference. The sequences used were chosen so as to represent the diversity through space, time, and host as well as possible.

Some of the branches connecting viruses with different hosts were long, perhaps reflecting a lack of sampling of the earliest viruses infecting the new host. This made the characterization of early adaptation events difficult, particularly for swine and canine lineages.

This work sheds light on the evolution of influenza viruses circulating in a variety of non-human hosts, which has not been studied as well as human influenza evolution. Higher dN/dS on the HA head in all mammalian hosts compared to avian host, and the retention or accumulation of large numbers of head glycosylations in some swine lineages (as occurs in human HA), suggest that mammals in general impose greater selection for antibody escape. Mammals also had higher dN/dS ratios in the stalk domain compared to birds. The reasons for this require further investigation, but it may reflect antigenic evolution in the stalk in mammals. In addition, evolutionary analysis of the sequences identified host adaptation in canines involving a particular phenotype. These results, based entirely on sequence data, provide information about influenza evolution in non-human hosts and place human influenza evolution in a larger context.

## MATERIALS AND METHODS

### Dataset Assembly

We downloaded all available hemagglutinin (HA) sequences for influenza A H3 and H1 from Genbank and downloaded specific geographical isolates, not available in Genbank, from GISAID on December 8th 2018. The sequences were aligned using the MAFFT [63] multiple sequence alignment tool. The aligned sequences were manually edited and cleaned in AliView version 1.26 software [64]. The resulting dataset was trimmed at the 5’ and 3’ ends to include solely the coding sequence. This dataset was subjected to multiple iterations of phylogeny reconstruction using FastTree version 1.0 [65] with a general time-reversible (GTR) model, and exclusion of outlier sequences whose genetic divergence and sampling date were incongruent using TempEst [66].

We defined lineages from the same host and geographical region, and only considered those circulating for at least three years in the region (Supplementary Table 1). Besides the H1 pandemic lineage circulating in humans since 2009, we defined 2 human H1 lineages circulating during the periods of 1918-1957 and 1977-2009. The H1 lineage reintroduced in 1977 derives directly from the viruses circulating in 1957, so studying the two lineages together would disrupt the temporal signal and consequently the molecular clock model, due to the lack of accumulation of mutations during the 20-year period where H1 did not circulate. For the non-avian lineages, we excluded sequences with mixed subtypes, as marked by “Hx” or “mixed”.

The number of sequences available makes a full Bayesian inference using all of them impractical due to the computations required. For this reason we analyzed a representative subset, which excluded those that cover less than 75% of the coding sequence with unambiguous bases, and retained only one of identical sequences collected in the same country on the same day, or in the same month or year in the case of incomplete date information.

Next, we applied a subsampling strategy that selected a specific number of sequences per country per time, in order to obtain similarly sized lineages that represented the overall circulating diversity. When sequences had the same country and collection date, we chose, in decreasing order of priority, those that were longer, had at least the month of collection rather than just the year, those with a smaller number of internal gaps with lengths not divisible by 3 (codons), and those with full dates of collection. We combined all subsampled lineages for each subtype and performed a final iteration of phylogeny reconstruction and exclusion of outlier sequences as described above, resulting in datasets with 873 sequences (16 lineages) for H3 and 964 sequences (33 lineages) for H1 (Figs. S1 and S2, and Files S1 and S2).

### Phylogenetic Analysis

Phylogenetic relationships were inferred separately for the H3 and H1 datasets, with a Bayesian phylogenetic approach using Markov chain Monte Carlo (MCMC) available via the BEAST v1.10.5 package [67] and the high-performance computational capabilities of the Biowulf Linux cluster at the National Institutes of Health, Bethesda, MD, USA (http://biowulf.nih.gov).

We divided the sequences into lineages based on host, geography, and a preliminary phylogeny inferred using FastTree with a GTR substitution model [68], retaining clusters with a minimum of 10 sequences and a minimum of three different collection years . Some lineages are paraphyletic with respect to other lineages. We partitioned the coding sequences into first, second and third codon positions and applied a separate Hasegawa-Kishino-Yano 85 (HKY85 [69]) substitution model with gamma-distributed rate variation among sites to all partitions [70]. We used an uncorrelated relaxed molecular clock with branch rates drawn from a lognormal distribution to account for evolutionary rate variation, and specified a Bayesian Skygrid coalescent tree prior [71]. For sequences with incomplete date of viral collection available, the lack of tip date precision was accommodated by sampling uniformly across a one-year window from January 1 to December 31.

We applied an approach that relies on counting methods, set in a Bayesian framework, to estimate lineage-specific synonymous (synonymous substitutions / all sites / year), non-synonymous (non-synonymous substitutions / all sites / year) substitution rates as well as their ratio (dN/dS or *ω*) in large datasets, while integrating over the posterior distribution of phylogenies and ancestral reconstructions to quantify the uncertainty on the lineage-specific estimates.

The nonsynonymous:synonymous ratios that we report are not, strictly speaking, dN/dS (*ω*), and the sequence changes are not properly referred to as substitutions, because many of the changes have not in any sense gone to fixation, and never will. We nevertheless continue a looser use of these terms that has become common in similar contexts. Terminology aside, these changes may have characteristics that differ from those that reflect the full effects of selection because they have come to predominate in their lineages.

The MCMC chain was run separately at least twenty times for each of the datasets and for at least 50 million iterations with variables recorded every 50,000 iterations, using the BEAGLE [72] library to improve computational performance. All parameters reached convergence, as assessed visually using Tracer v.1.7.1, with statistical uncertainty reflected in values of the 95% highest posterior density (HPD). At least 10% of the chain was removed from the beginning as burn-in. Maximum clade credibility (MCC) trees were summarized using TreeAnnotator v1.10.5 and visualized in FigTree v1.4.4.

We summarized the synonymous and nonsynonymous rate and dN/dS estimates for all protein sites (AS), head and stalk domains [73], and a set of 131 positions claimed to constitute the antigenically relevant sites for H3 (ARS, analyzed only for H3) [74]. All sites investigated are presented in Supplementary Table 2. We evaluated the statistical significance of differences in dN/dS by applying Fisher’s exact test to the counts of synonymous and nonsynonymous changes in each pair of lineages inferred from a most parsimonious reconstruction on the MCC tree. We then applied the Benjamini-Hochberg Procedure to find the set of pairwise comparisons giving a false discovery rate (FDR) of 5%.

### Ancestral Sequence Reconstruction

Ancestral state reconstruction of nucleotides was performed on the MCC tree using maximum parsimony, implemented as a Python program. In cases where the reconstruction was ambiguous, it was disambiguated arbitrarily for simplicity. Changes to the protein sequence were inferred from translations of the reconstructed nucleotide states.

### Glycosylation Patterns

We scanned all sequences (ancestral and contemporaneous) for each dataset to identify potential N-linked glycosylated sites, based on the motif Asn-X-Ser/Thr-X (N-X-S/T-X), where X is any amino acid other than proline (P) [75] using a program written in Python. We then plotted the number of glycosylated sites over time for each lineage and estimated their rate of accumulation or loss.

### Interface Residues in Equine and Canine H3

Each Protein Data Bank (PDB) entry was loaded into RasMol [76]. The amino acid atoms in one protein chain (HA1 or HA2) within 5 Angstroms of any amino acid atom in another protein chain were selected, and the amino acids to which they belonged were considered interface residues. For entries with chains that were not equivalent because they were related by non-crystallographic symmetry, each chain in the asymmetric unit was evaluated. Where necessary because of crystallographic symmetry, PDB biological assemblies were used. Rasmol was also used to produce the figures showing HA structures.

## ACKNOWLEDGMENTS

We thank Caitlin Bowers at Dartmouth College for help with compilation of the genetic datasets, and Emma Wagner at Dartmouth College and James R. Otieno at NIH/FIC for assistance with making figures in R. The opinions expressed in this article are those of the authors and do not reflect the view of the National Institutes of Health, the Department of Health and Human Services, or the United States government.

## Author contributions

Conceptualization: NST, MIN, JLC; Methodology: NST, PL, MIN, JLC; Investigation: SMK, NST, MIN, JLC; Visualization: NST, MIN, JLC; Supervision: PL, MIN, JLC; Writing - original draft: NST, MIN, JLC; Writing - review & editing: NST, PL, MIN, JLC

## Data and materials availability

No new data were generated or analyzed in support of this research. The XML-format files containing the data and model parametrization will be made available on GitHub (url will be added upon revision).

## REFERENCES

1. Johnson KEE, Song T, Greenbaum B, Ghedin E. Getting the flu: 5 key facts about influenza virus evolution. PLoS Pathog. 2017;13. doi:10.1371/journal.ppat.1006450

2. Widdowson M-A, Bresee JS, Jernigan DB. The Global Threat of Animal Influenza Viruses of Zoonotic Concern: Then and Now. J Infect Dis. 2017;216: S493–S498. doi:10.1093/infdis/jix331

3. Mostafa A, Abdelwhab EM, Mettenleiter TC, Pleschka S. Zoonotic Potential of Influenza A Viruses: A Comprehensive Overview. Viruses. 2018;10. doi:10.3390/v10090497

4. Gamblin SJ, Skehel JJ. Influenza Hemagglutinin and Neuraminidase Membrane Glycoproteins. J Biol Chem. 2010;285: 28403. doi:10.1074/jbc.R110.129809

5. Webster RG, Bean WJ, Gorman OT, Chambers TM, Kawaoka Y. Evolution and ecology of influenza A viruses. Microbiol Rev. 1992. pp. 152–79.

6. Song D, Kang B, Lee C, Jung K, Ha G, Kang D, et al. Transmission of Avian Influenza Virus (H3N2) to Dogs. Emerg Infect Dis. 2008;14: 741. doi:10.3201/eid1405.071471

7. Parrish CR, Murcia PR, Holmes EC. Influenza Virus Reservoirs and Intermediate Hosts: Dogs, Horses, and New Possibilities for Influenza Virus Exposure of Humans. J Virol. 2014 [cited 28 Jan 2022]. Available: https://journals.asm.org/doi/abs/10.1128/JVI.03146-14

8. Voorhees IEH, Dalziel BD, Glaser A, Dubovi EJ, Murcia PR, Newbury S, et al. Multiple Incursions and Recurrent Epidemic Fade-Out of H3N2 Canine Influenza A Virus in the United States. J Virol. 2018 [cited 28 Jan 2022]. doi:10.1128/JVI.00323-18

9. Guan L, Shi J, Kong X, Ma S, Zhang Y, Yin X, et al. H3N2 avian influenza viruses detected in live poultry markets in China bind to human-type receptors and transmit in guinea pigs and ferrets. Emerg Microbes Infect. 2019;8: 1280. doi:10.1080/22221751.2019.1660590

10. Trovão NS, Nelson MI. When Pigs Fly: Pandemic influenza enters the 21st century. PLoS Pathog. 2020;16. doi:10.1371/journal.ppat.1008259

11. Olguin-Perglione C, Barrandeguy ME. An Overview of Equine Influenza in South America. Viruses. 2021;13: 888. doi:10.3390/v13050888

12. World Health Organization. Influenza (Seasonal) Fact sheet. 2018 [cited 28 Jan 2022]. Available: https://www.who.int/news-room/fact-sheets/detail/influenza-(seasonal)

13. Boni MF, Galvani AP, Wickelgren AL, Malani A. Economic epidemiology of avian influenza on smallholder poultry farms. Theor Popul Biol. 2013;90: 135. doi:10.1016/j.tpb.2013.10.001

14. Hill EM, House T, Dhingra MS, Kalpravidh W, Morzaria S, Osmani MG, et al. The impact of surveillance and control on highly pathogenic avian influenza outbreaks in poultry in Dhaka division, Bangladesh. PLoS Comput Biol. 2018;14. doi:10.1371/journal.pcbi.1006439

15. Kemen MJ, Frank RA, Babish JB. An outbreak of equine influenza at a harness horse racetrack. Cornell Vet. 1985;75: 277–288.

16. Sack A, Cullinane A, Daramragchaa U, Chuluunbaatar M, Gonchigoo B, Gray GC. Equine Influenza Virus—A Neglected, Reemergent Disease Threat. Emerg Infect Dis. 2019;25: 1185. doi:10.3201/eid2506.161846

17. Cullinane A, Gahan J, Walsh C, Nemoto M, Entenfellner J, Olguin-Perglione C, et al. Evaluation of Current Equine Influenza Vaccination Protocols Prior to Shipment, Guided by OIE Standards. Vaccines. 2020;8. doi:10.3390/vaccines8010107

18. Taubenberger JK, M MD. 1918 Influenza: the mother of all pandemics. Emerg Infect Dis. 2006. doi:10.3201/eid1209.05-0979

19. Jester BJ, Uyeki TM, Jernigan DB. Fifty Years of Influenza A(H3N2) Following the Pandemic of 1968. Am J Public Health. 2020;110: 669. doi:10.2105/AJPH.2019.305557

20. Murcia PR, Wood JLN, Holmes EC. Genome-Scale Evolution and Phylodynamics of Equine H3N8 Influenza A Virus. J Virol. 2011;85: 5312. doi:10.1128/JVI.02619-10

21. Anderson TK, Macken CA, Lewis NS, Scheuermann RH, Reeth KV, Brown IH, et al. A Phylogeny-Based Global Nomenclature System and Automated Annotation Tool for H1 Hemagglutinin Genes from Swine Influenza A Viruses. mSphere. 2016;1. doi:10.1128/mSphere.00275-16

22. Collins PJ, Vachieri SG, Haire LF, Ogrodowicz RW, Martin SR, Walker PA, et al. Recent evolution of equine influenza and the origin of canine influenza. Proc Natl Acad Sci U S A. 2014;111: 11175–11180. doi:10.1073/pnas.1406606111

23. Mena I, Nelson MI, Quezada-Monroy F, Dutta J, Cortes-Fernandez R, Lara-Puente JH, et al. Origins of the 2009 H1N1 influenza pandemic in swine in Mexico. Elife. 2016. doi:10.7554/eLife.16777

24. Plotkin JB, Kudla G. Synonymous but not the same: the causes and consequences of codon bias. Nat Rev Genet. 2011;12: 32. doi:10.1038/nrg2899

25. Sandt CE van de, Kreijtz JHCM, Rimmelzwaan GF. Evasion of Influenza A Viruses from Innate and Adaptive Immune Responses. Viruses. 2012;4: 1438. doi:10.3390/v4091438

26. Shi Y, Wu Y, Zhang W, Qi J, Gao GF. Enabling the “host jump”: structural determinants of receptor-binding specificity in influenza A viruses. Nat Rev Microbiol. 2014;12: 822–831. doi:10.1038/nrmicro3362

27. Bouvier NM, Palese P. The biology of influenza viruses. Vaccine. 2008;26 Suppl 4: D49–53. doi:10.1016/j.vaccine.2008.07.039

28. Shaw M, Palese P. Fields virology, p 1151–1185. Fields Virol 6th Ed Lippincott Williams Wilkins Phila PA. 2013.

29. Kirkpatrick E, Qiu X, Wilson PC, Bahl J, Krammer F. The influenza virus hemagglutinin head evolves faster than the stalk domain. Sci Rep. 2018;8: 10432. doi:10.1038/s41598-018-28706-1

30. Altman MO, Angel M, Košík I, Trovão NS, Zost SJ, Gibbs JS, et al. Human Influenza A Virus Hemagglutinin Glycan Evolution Follows a Temporal Pattern to a Glycan Limit. mBio. 2019;10. doi:10.1128/mBio.00204-19

31. Reading PC, Tate MD, Pickett DL, Brooks AG. Glycosylation as a target for recognition of influenza viruses by the innate immune system. Adv Exp Med Biol. 2007;598: 279–292. doi:10.1007/978-0-387-71767-8_20

32. Kosik I, Ince WL, Gentles LE, Oler AJ, Kosikova M, Angel M, et al. Influenza A virus hemagglutinin glycosylation compensates for antibody escape fitness costs. PLoS Pathog. 2018;14. doi:10.1371/journal.ppat.1006796

33. Watanabe Y, Bowden TA, Wilson IA, Crispin M. Exploitation of glycosylation in enveloped virus pathobiology. Biochim Biophys Acta Gen Subj. 2019;1863: 1480. doi:10.1016/j.bbagen.2019.05.012

34. Taubenberger JK, C KJ. Influenza virus evolution, host adaptation, and pandemic formation. Cell Host Microbe. 2010. doi:10.1016/j.chom.2010.05.009

35. Osterholm MT, Kelley NS, Manske JM, Ballering KS, Leighton TR, Moore KA. The Compelling Need for Game- Changing Influenza Vaccines. In: CIDRAP [Internet]. Oct 2012 [cited 28 Jan 2022]. Available: https://www.cidrap.umn.edu/compelling-need-game-changing-influenza-vaccines

36. Yamayoshi S, Kawaoka Y. Current and future influenza vaccines. Nat Med. 2019;25: 212– 220. doi:10.1038/s41591-018-0340-z

37. Van Reeth K, Ma W. Swine Influenza Virus Vaccines: To Change or Not to Change— That’s the Question. In: Richt JA, Webby RJ, editors. Swine Influenza. Berlin, Heidelberg: Springer; 2013. pp. 173–200. doi:10.1007/82_2012_266

38. Vincent AL, Perez DR, Rajao D, Anderson TK, Abente EJ, Walia RR, et al. Influenza A virus vaccines for swine. Vet Microbiol. 2017;206: 35–44. doi:10.1016/j.vetmic.2016.11.026

39. Rodriguez L, Nogales A, Reilly EC, Topham DJ, Murcia PR, Parrish CR, et al. A live-attenuated influenza vaccine for H3N2 canine influenza virus. Virology. 2017;504: 96–106. doi:10.1016/j.virol.2017.01.020

40. Chambers TM. A brief introduction to equine influenza and equine influenza viruses. Methods Mol Biol Clifton NJ. 2014;1161: 365–370. doi:10.1007/978-1-4939-0758-8_31

41. Tabynov K, Kydyrbayev Z, Ryskeldinova S, Assanzhanova N, Kozhamkulov Y, Inkarbekov D, et al. Safety and immunogenicity of a novel cold-adapted modified-live equine influenza virus vaccine. Aust Vet J. 2014;92: 450–457. doi:10.1111/avj.12248

42. Na W, Yeom M, Yuk H, Moon H, Kang B, Song D. Influenza virus vaccine for neglected hosts: horses and dogs. Clin Exp Vaccine Res. 2016;5: 117. doi:10.7774/cevr.2016.5.2.117

43. Swayne DE. Avian influenza vaccines and therapies for poultry. Comp Immunol Microbiol Infect Dis. 2009;32: 351–363. doi:10.1016/j.cimid.2008.01.006

44. Nelson MI, Holmes EC. The evolution of epidemic influenza. Nat Rev Genet. 2007;8: 196– 205. doi:10.1038/nrg2053

45. Yoo SJ, Kwon T, Lyoo YS. Challenges of influenza A viruses in humans and animals and current animal vaccines as an effective control measure. Clin Exp Vaccine Res. 2018;7: 1. doi:10.7774/cevr.2018.7.1.1

46. Sautto GA, Kirchenbaum GA, Ross TM. Towards a universal influenza vaccine: different approaches for one goal. Virol J. 2018;15: 17. doi:10.1186/s12985-017-0918-y

47. Bedford T, A SM, Philippe L, Gytis D, Victoria G, J HA, et al. Integrating influenza antigenic dynamics with molecular evolution. Elife. 2014. p. e01914. doi:10.7554/eLife.01914

48. Lewis NS, A RC, Pinky L, K AT, Kathryn B, Filip B, et al. The global antigenic diversity of swine influenza A viruses. eLife. 2016. doi:10.7554/eLife.12217

49. Park J-K, Xiao Y, Ramuta MD, Rosas LA, Fong S, Matthews AM, et al. Pre-existing immunity to influenza virus hemagglutinin stalk might drive selection for antibody-escape mutant viruses in a human challenge model. Nat Med. 2020;26: 1240–1246. doi:10.1038/s41591-020-0937-x

50. Wrigley NG, Brown EB, Daniels RS, Douglas AR, Skehel JJ, Wiley DC. Electron microscopy of influenza haemagglutinin-monoclonal antibody complexes. Virology. 1983;131: 308–314. doi:10.1016/0042-6822(83)90499-3

51. Skehel JJ, Stevens DJ, Daniels RS, Douglas AR, Knossow M, Wilson IA, et al. A carbohydrate side chain on hemagglutinins of Hong Kong influenza viruses inhibits recognition by a monoclonal antibody. Proc Natl Acad Sci U S A. 1984;81: 1779–1783. doi:10.1073/pnas.81.6.1779

52. Abe Y, Takashita E, Sugawara K, Matsuzaki Y, Muraki Y, Hongo S. Effect of the addition of oligosaccharides on the biological activities and antigenicity of influenza A/H3N2 virus hemagglutinin. J Virol. 2004;78: 9605–9611. doi:10.1128/JVI.78.18.9605-9611.2004

53. Gallagher P, Henneberry J, Wilson I, Sambrook J, Gething MJ. Addition of carbohydrate side chains at novel sites on influenza virus hemagglutinin can modulate the folding, transport, and activity of the molecule. J Cell Biol. 1988;107: 2059–2073. doi:10.1083/jcb.107.6.2059

54. Sun S, Wang Q, Zhao F, Chen W, Li Z. Glycosylation Site Alteration in the Evolution of Influenza A (H1N1) Viruses. PLOS ONE. 2011;6: e22844. doi:10.1371/journal.pone.0022844

55. Mancera Gracia JC, Pearce DS, Masic A, Balasch M. Influenza A Virus in Swine: Epidemiology, Challenges and Vaccination Strategies. Front Vet Sci. 2020;7: 647. doi:10.3389/fvets.2020.00647

56. Kong W, Ye J, Guan S, Liu J, Pu J. Epidemic status of Swine influenza virus in china. Indian J Microbiol. 2014;54: 3–11. doi:10.1007/s12088-013-0419-7

57. Wen F, Blackmon S, Olivier AK, Li L, Guan M, Sun H, et al. Mutation W222L at the Receptor Binding Site of Hemagglutinin Could Facilitate Viral Adaption from Equine Influenza A(H3N8) Virus to Dogs. J Virol. 2018;92: e01115–18. doi:10.1128/JVI.01115-18

58. Russell CJ. Acid-induced membrane fusion by the hemagglutinin protein and its role in influenza virus biology. Curr Top Microbiol Immunol. 2014;385: 93–116. doi:10.1007/82_2014_393

59. Galloway SE, Reed ML, Russell CJ, Steinhauer DA. Influenza HA subtypes demonstrate divergent phenotypes for cleavage activation and pH of fusion: implications for host range and adaptation. PLoS Pathog. 2013;9: e1003151. doi:10.1371/journal.ppat.1003151

60. Mair CM, Ludwig K, Herrmann A, Sieben C. Receptor binding and pH stability - how influenza A virus hemagglutinin affects host-specific virus infection. Biochim Biophys Acta. 2014;1838: 1153–1168. doi:10.1016/j.bbamem.2013.10.004

61. Byrd-Leotis L, Galloway SE, Agbogu E, Steinhauer DA. Influenza hemagglutinin (HA) stem region mutations that stabilize or destabilize the structure of multiple HA subtypes. J Virol. 2015;89: 4504–4516. doi:10.1128/JVI.00057-15

62. Di Lella S, Herrmann A, Mair CM. Modulation of the pH Stability of Influenza Virus Hemagglutinin: A Host Cell Adaptation Strategy. Biophys J. 2016;110: 2293–2301. doi:10.1016/j.bpj.2016.04.035

63. Katoh K, Toh H. Recent developments in the MAFFT multiple sequence alignment program. Brief Bioinform. 2008. pp. 286–98. doi:10.1093/bib/bbn013

64. Larsson A. AliView: a fast and lightweight alignment viewer and editor for large datasets. Bioinforma Oxf Engl. 2014;30: 3276–3278. doi:10.1093/bioinformatics/btu531

65. Price MN, Dehal PS, Arkin AP. FastTree: computing large minimum evolution trees with profiles instead of a distance matrix. Mol Biol Evol. 2009;26: 1641–1650. doi:10.1093/molbev/msp077

66. Rambaut A, Lam TT, Max Carvalho L, Pybus OG. Exploring the temporal structure of heterochronous sequences using TempEst (formerly Path-O-Gen). Virus Evol. 2016. p. vew007. doi:10.1093/ve/vew007

67. Suchard MA, Lemey P, Baele G, Ayres DL, Drummond AJ, Rambaut A. Bayesian phylogenetic and phylodynamic data integration using BEAST 1.10. Virus Evol. 2018;4: vey016. doi:10.1093/ve/vey016

68. Price MN, Dehal PS, Arkin AP. FastTree 2 – Approximately Maximum-Likelihood Trees for Large Alignments. PLoS ONE. 2010;5. doi:10.1371/journal.pone.0009490

69. Hasegawa M, Kishino H, Yano T. Dating of the human-ape splitting by a molecular clock of mitochondrial DNA. J Mol Evol. 1985;22: 160–174. doi:10.1007/BF02101694

70. Shapiro B, Rambaut A, Drummond AJ. Choosing appropriate substitution models for the phylogenetic analysis of protein-coding sequences. Mol Biol Evol. 2006;23: 7–9. doi:10.1093/molbev/msj021

71. Gill MS, Lemey P, Faria NR, Rambaut A, Shapiro B, Suchard MA. Improving Bayesian population dynamics inference: a coalescent-based model for multiple loci. Mol Biol Evol. 2013;30: 713–724. doi:10.1093/molbev/mss265

72. Ayres DL, Cummings MP, Baele G, Darling AE, Lewis PO, Swofford DL, et al. BEAGLE 3: Improved Performance, Scaling, and Usability for a High-Performance Computing Library for Statistical Phylogenetics. Syst Biol. 2019;68: 1052–1061. doi:10.1093/sysbio/syz020

73. Krammer F, Palese P. Influenza virus hemagglutinin stalk-based antibodies and vaccines. Curr Opin Virol. 2013;3: 521–530. doi:10.1016/j.coviro.2013.07.007

74. Bush RM, Bender CA, Subbarao K, Cox NJ, Fitch WM. Predicting the evolution of human influenza A. Science. 1999;286: 1921–1925. doi:10.1126/science.286.5446.1921

75. Mellquist JL, Kasturi L, Spitalnik SL, Shakin-Eshleman SH. The amino acid following an asn-X-Ser/Thr sequon is an important determinant of N-linked core glycosylation efficiency. Biochemistry. 1998;37: 6833–6837. doi:10.1021/bi972217k

76. Sayle RA, Milner-White EJ. RASMOL: biomolecular graphics for all. Trends Biochem Sci. 1995;20: 374. doi:10.1016/s0968-0004(00)89080-5

